# Delineating the heterogeneity of matrix-directed differentiation towards soft and stiff tissue lineages via single-cell profiling

**DOI:** 10.1101/2020.11.23.394460

**Authors:** Shlomi Brielle, Danny Bavli, Alex Motzik, Yoav Kan-Tor, Batia Avni, Oren Ram, Amnon Buxboim

## Abstract

Mesenchymal stromal/stem cells (MSCs) are a heterogeneous population of multipotent progenitors that contribute to tissue regeneration and homeostasis. MSCs assess extracellular elasticity by probing resistance to applied forces via adhesion, cytoskeletal, and nuclear mechanotransducers, that direct differentiation toward soft or stiff tissue lineages. Even under controlled conditions, MSC differentiation exhibits substantial cell-to-cell variation that remains poorly characterized. By single-cell transcriptional profiling of naïve, matrix-conditioned, and early differentiation state cells, we identified distinct MSC subpopulations with distinct mechanosensitivities, differentiation capacities, and cell cycling. We showed that soft matrices support adipogenesis of multipotent cells and endochondral ossification of non-adipogenic cells, whereas intramembranous ossification and pre-osteoblast proliferation are enhanced by stiff matrices. Using diffusion pseudotime mapping, we delineated hierarchical matrix-directed differentiation and identified mechanoresponsive genes. We found that tropomyosin-1 (*TPM1*) is highly sensitive to stiffness cues both at RNA and protein levels and that changes in expression of TPM1 determine adipogenic or osteogenic fates. Thus, cell-to-cell variation in tropomyosin-mediated matrix-sensing contributes to impaired differentiation with implications to the biomedical potential of MSCs.

## Main

MSCs are present in all vascularized compartments owing to their perivascular origin, and as such they can be isolated from bone marrow, fat, placenta, and other tissues.^1,2^ The definition of MSCs relies on their surface-adherence and expansion in culture, the expression of several mesodermal and absence of hematopoietic surface markers, and retaining a multilineage differentiation capacity towards fat, cartilage and bone under defined induction media. This elusive definition permits a significant molecular and phenotypic variation between cells that were derived from different donors, different tissues of the same donor, different clones isolated from the same tissue, and different cells of the same clone.^3–5^ MSCs have drawn considerable clinical interest for mediating immunomodulatory effects rather than for their “stemness”.^6^ However, the therapeutic potential of MSC-based treatments is compromised by their heterogeneous nature by diluting clinically-functional cells and by contributing to the inconsistent clinical outcomes across treatments.^7,8^ Characterizing MSC heterogeneity and its clinical implications will thus improve experimental reproducibility and biomedical standardization.

MSCs are highly sensitive to the mechanical properties of their microenvironment. These extracellular, tissue-specific cues are actively probed by all adherent cells, ^9–12^ whereas impaired mechanosensitivity is leveraged by oncogenically transformed cells for evading apoptotic pathways.^13^ The mechanical resistance of the cellular microenvironment to cell-generated forces is set by extracellular elasticity and geometrical boundary conditions.^14^ These stress-strain relationships can be converted into biochemical signals through the forced unfolding of linker proteins,^15,16^ force-sensitive^17^ and catch-bond adhesions to extracellular matrix ligands^18,19^ and to neighboring cell receptors,^20,21^ tension-mediated filament-stabilization,^22,23^ or direct physical stretching of chromatin loci in the nucleus.^24^ The emerging intracellular signals are mediated via a number of pathways that regulate gene expression and direct cell-fate decisions.^25,26^ The resulting upregulation of cytoskeletal and force-generating target genes stabilizes a contractile cell state with positive feedback to extracellular stiffness.^27,28^

Here, we exposed naïve bone-marrow derived clinically-graded MSCs, which had been harvested for bone-marrow transplantation treatments, to matrices with controlled elasticities and to a bipotential induction cocktail that permits differentiation towards fat or bone. To gain insight into the implications of MSC heterogeneity on cellular mechanosensitivity and multipotency, we transcriptionally profiled the cells via whole genome single-cell RNA sequencing at naïve, matrix conditioned, and early differentiating stages. Unsupervised clustering of MSC subpopulations and diffusion pseudotime mapping revealed a bifurcation of cell state propagation between differentiated and non-differentiated fates. Whole genome screening highlighted tropomyosin-1 as a matrix-responsive gene, which was experimentally validated. Using targeted gene silencing and overexpression, tropomyosin-1 was found to be a highly potent regulator of cell differentiation downstream of tissue-level matrix mechanics. Characterizing cell-to-cell variations among the response to matrix and differentiation cues during cell-state propagation contributes to elucidating MSC heterogeneity with future implications to cell-based therapeutics.

## Results

### Cell-to-cell variation in MSC mechanosensitivity

MSC differentiation toward soft and stiff tissue lineages is a tightly regulated process that integrates mechanical inputs and biochemical cues from the microenvironments.^29–31^ Here we studied low-passage, naïve MSCs that were obtained from bone marrow donors during allogeneic transplantation. To study how matrix elasticity directs differentiation toward adipogenesis or osteogenesis leading to fat and bone lineages, respectively, we expanded naïve MSCs on polystyrene and seeded them on collagen-coated hydrogel substrates with controlled stiffness: The “soft” collagen-coated hydrogel matrix (2 kPa) mimics the elasticity of adipose tissue^32^ and the “stiff” chondrogenic peri-cellular matrix and osteoid matrix (25 kPa)^29,33,34^ mimics the cartilaginous endochondral ossification/osteoid microenvironment (Fig. 1a-i).^14^ Cells were matrix-conditioned in basal medium for 3 days before basal medium was replaced with bi-potential induction medium that permits adipogenic and osteogenic differentiation (Fig. 1a-ii).^35,36^ MSCs cultured on stiff matrix appear to spread more than those cultured on soft matrix (Fig 1b-i); the stiff matrix provides support for the striated organization of mature actomyosin stress fibers.^37^ Cell and nucleus projected areas are established markers of cell mechanosensitivity, yet only 25% of the cells spread more and nuclei became more stretched and flattened on stiff matrices than soft, thus reflecting cellular heterogeneity (Fig. 1b-ii). Adipogenic differentiation was favored on soft matrices (Fig. S1a), and osteogenic differentiation was favored on stiff matrices (Fig. S1b). Nevertheless, most cells failed to undergo adipogenesis even under optimal adipogenic conditions, and a fraction of cells differentiated counter to matrix elasticity. Thus, population averages of matrix-directed cytoskeletal organization, cell and nucleus projected morphologies, and cell differentiation confirmed active mechanosensitivity^9^, but the observed cell-to-cell variability is indicative of a heterogeneous response to mechanical cues as previously reported.^4,5,38^

**Figure 1.**
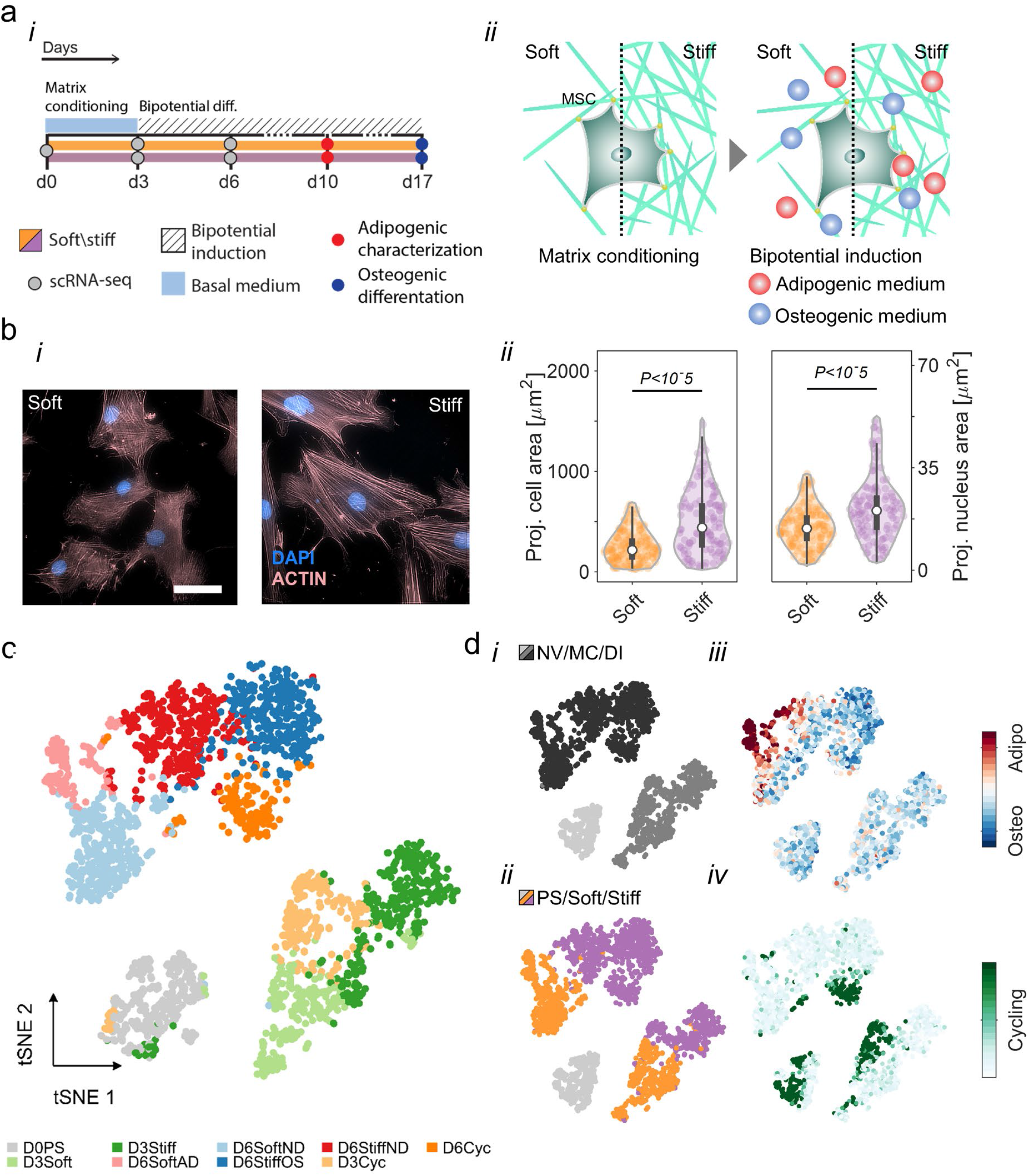
Resolving MSC heterogeneity using matrix elasticity and bi-potential differentiation induction signaling. **a,** (i) Experimental design: Naïve MSCs were cultured on soft (2 kPa) and stiff (25 kPa) collagen-coated hydrogel matrices that mimic fat and that mimic the cartilaginous endochondral ossification/osteoid microenvironment, respectively. (ii) Cells were matrix-conditioned for 3 days in basal medium, followed by a switch to bi-potential induction medium for 3 days. Adipogenic and osteogenic differentiation potentials were evaluated on day 10 and day 17, respectively. Naïve, matrix conditioned, and early differentiating MSCs were harvested on day 0 (388 cells), day 3 (soft: 467 cells; stiff: 450 cells) and day 6 (soft: 534 cells; stiff: 951 cells), respectively, and analyzed by single-cell transcriptional profiling. **b,** (i) Phalloidin staining of MSCs cultured on soft matrices (left) and on stiff matrices (right) show striated organization of mature actomyosin stress fibers on stiff. Scale bar, 50 μm. (ii) Average cell and nucleus projected areas on soft and stuff matrices. **c,** Single-cell transcriptomes were divided into nine subpopulations using unsupervised *k*-means clustering at the dimensionality reduced principal component space. Associations between subpopulations were projected onto a t-SNE map. **d,** MSC subpopulations are characterized by (i) cell state (ii) matrix elasticity, (iii) early differentiation, and (iv) cell cycling. NV: naïve. MC: matrix conditioning. DI: differentiation induction.

### Matrix sensitivity, early differentiation, and cell cycling define distinctive MSC subpopulations

To characterize cell-to-cell variation in matrix-directed cell-fate decisions, we employed microfluidics-based single-cell RNA sequencing^39,40^ and profiled transcriptomes of cells in the naïve state (388 cells), matrix-conditioned state (soft: 467 cells, stiff: 450 cells), and post-differentiation induction (soft: 534, stiff: 951 cells). Single-cell transcriptomes were dimensionally reduced via principal component analysis (PCA) of highly variable genes (Fig. S2a). Transcriptomes of matrix-conditioned and post-differentiation induction cells formed clusters that were defined by cell state (naïve, matrix conditioned, and early differentiation) and matrix elasticity (Fig. S2b). All single-cell transcriptomes were divided into nine subpopulations using unsupervised *k*-means clustering in the PCA space and projected onto a t-distributed stochastic neighbor embedding (t-SNE) map (Fig. 1c). Matrix-conditioned cells analyzed on day 3 were divided between soft and stiff clusters (D3Soft and D3Stiff) and a cohort of cycling cells that were cultured on both matrices (D3Cyc). Early differentiation state cells, analyzed on day 6, were divided between adipogenic (D6SoftAD) and non-differentiating (D6SoftND) soft matrix clusters and osteogenic (D6StiffOS) and non-differentiating (D6StiffND) stiff matrix clusters. Unsupervised clustering paralleled the experimental parameters cell state and matrix elasticity (Fig. 1d-i and 1d-ii) and correlated with established gene signatures of adipogenic^41,42^ and osteogenic^41,43–46^ differentiation and cell cycling^40^ (Fig. 1d-iii and 1d-iv; Table-1). Unlike matrix-conditioned cycling subpopulation analyzed on day 3, the post-differentiation induction cycling population analyzed on day 6 was enriched for osteogenic cells that were cultured on stiff matrices. Indeed, pre-osteoblast proliferation is consistent with direct intramembranous ossfication.^47,48^

**Table 1.**
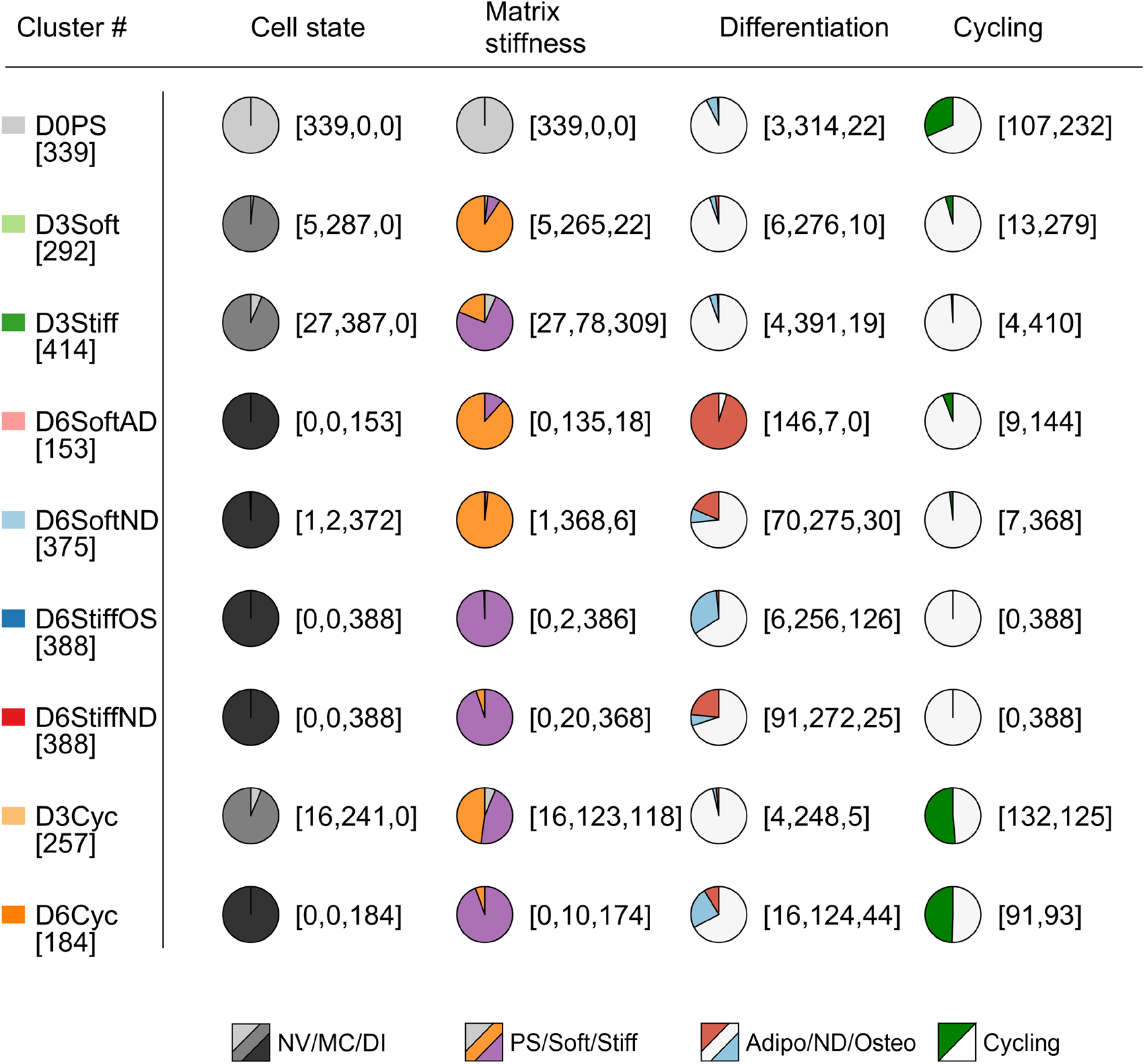
MSC subpopulations are defined by cell state, matrix elasticity, differentiation induction, and cell cycling. Subpopulations of single-cell transcriptomes are characterized by enrichment of day of culture, matrix elasticity, early differentiation, and cell cycling. Cell numbers are specified in brackets. NV: naïve. MC: matrix conditioning. DI: differentiation induction.

Clustering of gene intensities across single-cell transcriptomes highlights the associations of naive, matrix conditioned, and early differentiation state subpopulations (Fig. 2a). Genes that were differentially expressed across cell states (Fig. 2b-i) underlie the transition from polystyrene (naïve) to collagen-coated matrices (matrix conditioning) to bi-potential induction medium (early differentiation). Extracellular matrix (ECM) and cell adhesion genes encoding paxillin, elastin, and aggrecan were upregulated in naïve cells, whereas genes encoding vinculin, fibronectin-1, and lysyl oxidase collagen crosslinker were upregulated in matrix-conditioned cells. Matrix adhesion and ECM genes were downregulated in early differentiating cells, and insulin growth factors 1 and 2 and the related binding protein gene *IGFBP2* were upregulated (Fig. 2b-i). Next, we identified genes that were differentially expressed on soft versus stiff matrices (Fig. 2b-ii). Stiff matrices supported upregulation of genes involved in actin binding and the actomyosin cytoskeleton during matrix conditioning in basal medium (D3Stiff vs. D3Soft) and in non-osteogenic cells (D6StiffND vs. D6SoftAD) and non-adipogenic cells (D6StiffOS vs. D6SoftND) under bi-potential induction conditions.

**Figure 2.**
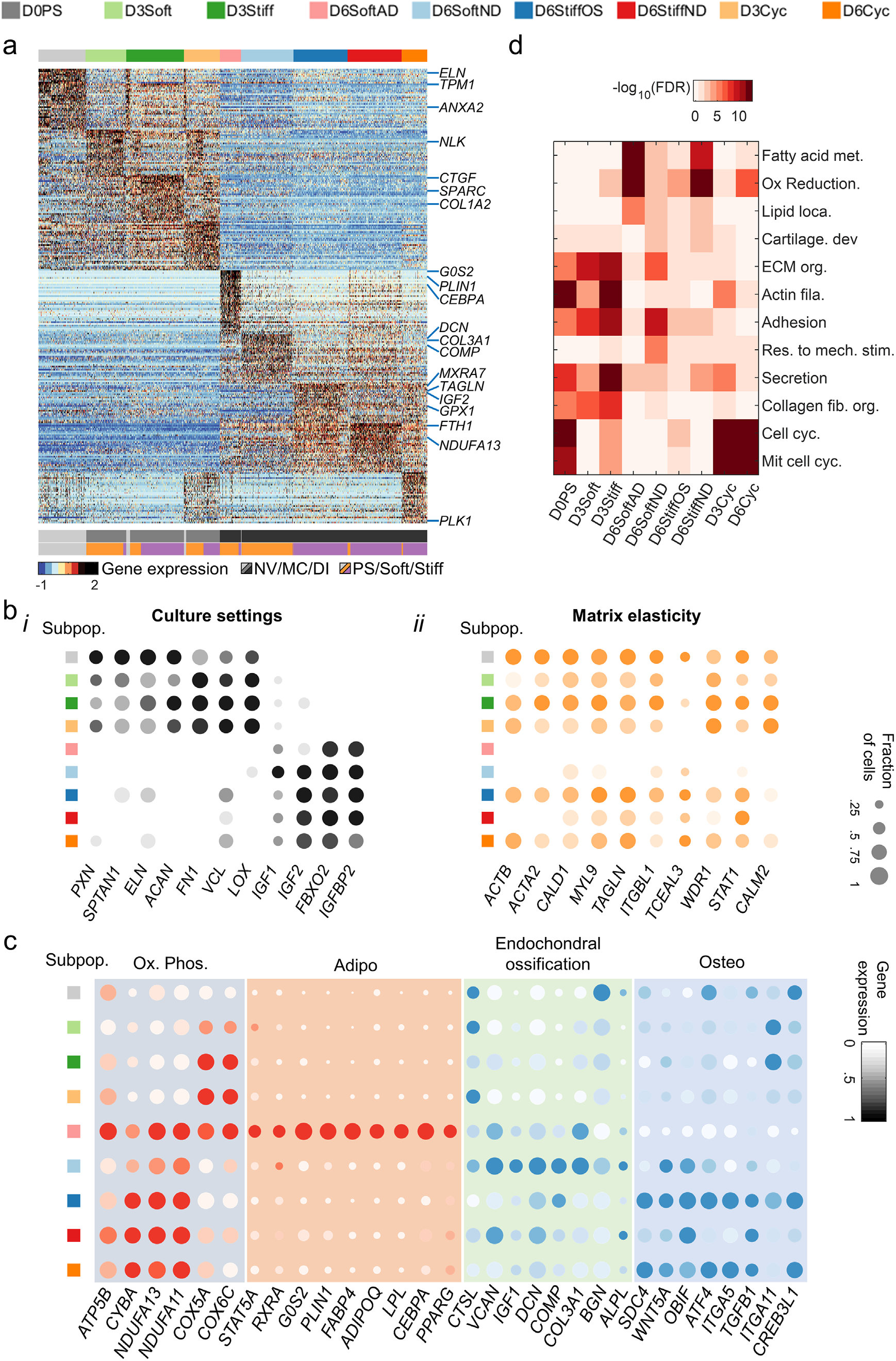
Differential gene expression analysis reveals cell-fate decisions of MSC subpopulations. **a,** Color-coded heatmap of z-score-normalized differentially expressed genes shows separation between naïve, matrix-conditioned, and early differentiating subpopulations (lfc > 1; adjusted *p*-value<10^−5^). **b** Average intensities and fraction of positive cells are plotted for genes that are differentially expressed between (i) culture settings and (ii) matrix elasticities. **c**, Day 6 subpopulations are characterized by expression patterns of OxPhos, adipogenesis, endochondral ossification, and osteogenesis. **d,** GO term analysis of differentially expressed genes reveals enrichment of metabolic, cell adhesion, cytoskeletal, ECM, differentiation, and cell-cycling patterns that characterize MSC subpopulations (Benjamini-Hochberg corrected). GO: Gene Ontology. ECM: extracellular matrix. OxPhos: oxidative phosphorylation.

On soft matrix, upregulation of oxidative phosphorylation genes, which are associated with biogenesis during pre-adipocyte differentiation,^49^ parallels upregulation of adipogenic markers in D6SoftAD cells that differentiate into adipocytes (Fig. 2c). In non-adipogenic cells that are cultured also on soft matrices (D6SoftND), nuclear-encoded oxidative phosphorylation genes are downregulated and endochondral ossification markers of indirect osteoblast differentiation are upregulated (Fig. 2c).^50^ Endochondral ossification is linked with a low oxidative phosphorylation state already during day-3 soft matrix conditioning (Fig. S3). Unlike soft matrices that support adipogenesis and endochondral ossification, direct osteoblast differentiation via membranous ossification (D6StiffOS and D6Cyc cells) is supported by stiff matrices as indicated by upregulation of osteogenic markers. Gene ontology (GO) analysis showed enrichment of genes annotated with fatty acid metabolism, oxidative reduction, and lipid localization terms in D6SoftAD cells; in cartilage organization in D6SoftND cells but not D6StiffOS cells; and in ECM, adhesion, and actin cytoskeletal terms in stiff-matrix (D3Stiff) but not soft-matrix conditioned cells (D3Soft) (Fig. 2d).

### Matrix-directed cell-fate decision-making processes revealed by diffusion pseudotime mapping

The single-cell transcriptomes provide multicellular snapshots of matrix-directed cell-fate decision making that highlight cell-to-cell variation. To reconstruct the effective propagation from a naïve cell state through matrix conditioning to early differentiation, we employed diffusion pseudotime analysis, which measures random-walk transcriptomic distances between cell states (Fig. 3a-b).^51,52^ Cells propagated from D0PS to D3Soft or D3Stiff states and bifurcated between adipogenic and non-differentiated fates on soft matrix. On stiff matrix, the bifurcation was between osteogenic and non-differentiated fates. Upstream of bifurcation, the pseudotime propagation rate appears to be slower on stiff matrix than soft matrix. Differences in pseudotime rates suggest that during matrix conditioning, soft matrices support induction of expression of adipogenic genes concomitantly with the suppression of osteogenic genes that are continuously expressed in naïve MSCs at low levels.^41^ On the soft matrix, cell cycling (D3Cyc) occurs concomitantly with matrix conditioning. In contrast, on the stiff matrix, cell cycling follows matrix conditioning and directs osteogenic differentiation. D6Cyc parallels D6StiffOS.

**Figure 3.**
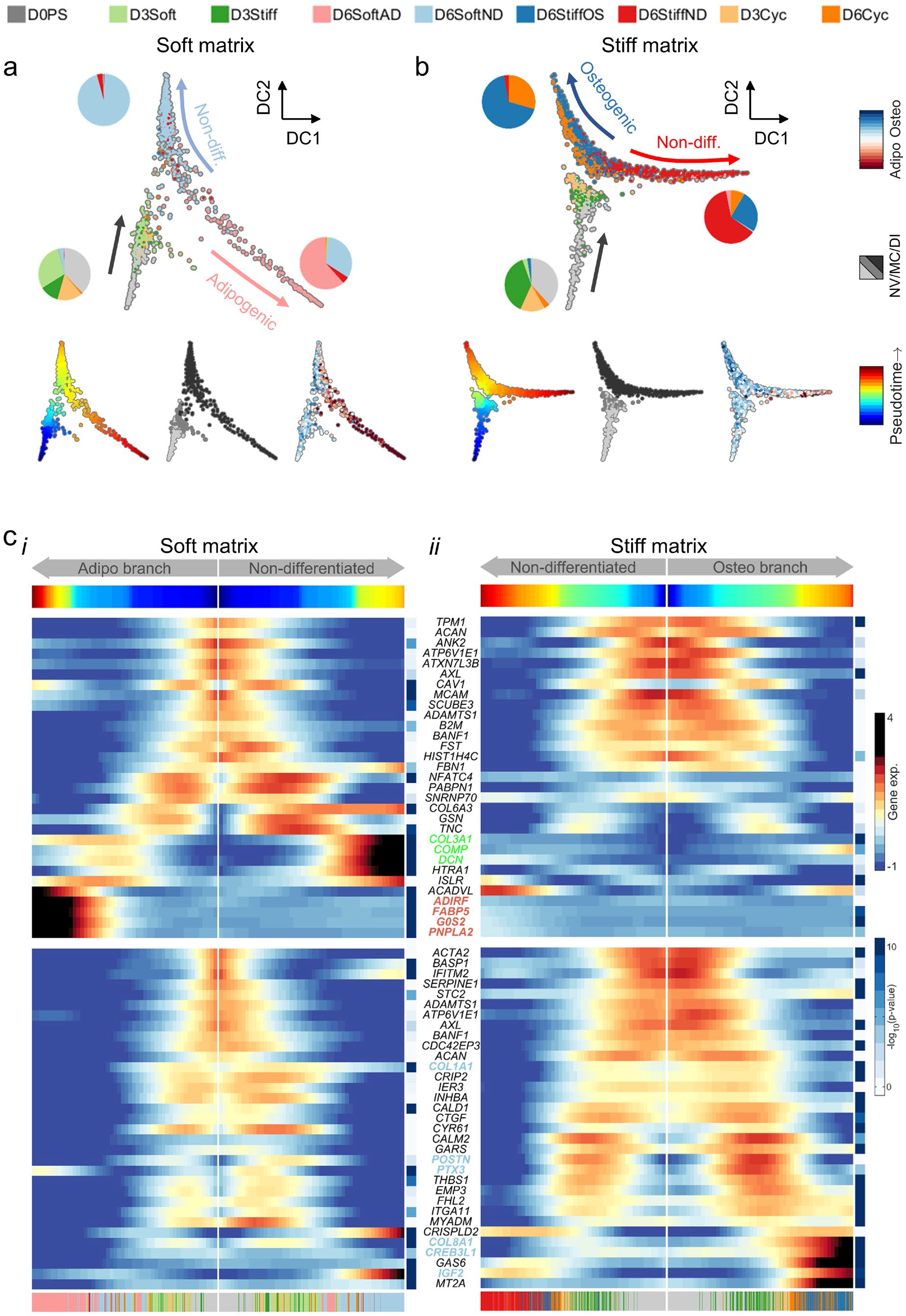
Diffusion pseudotime dynamics characterizes matrix-directed cell-fate decision-making. **a-b,** Diffusion mapping characterizes MSC propagation from naïve to matrix-conditioned states and bifurcation between (a) adipogenic and non-differentiated fates on soft matrix and (b) osteogenic and non-differentiated fates on stiff matrix. **c**, Pseudotime projections of differentially expressed genes (adjusted *p*-value < 10^−5^) that were upregulated in (i) cells cultured on soft matrix (top panels) and (ii) in cells cultured on stiff matrix. Endochondral ossification (green) and adipogenic (red) markers are upregulated along soft matrix branches, and osteogenic markers (blue) are upregulated on a stiff matrix branch. NV: naïve. MC: matrix conditioning. DI: differentiation induction.

We next used the principle elastic tree algorithm to evaluate the effective transcriptome date. The Y-shaped scaffold trees are divided into a naïve state and matrix conditioning branch, adipogenic (soft matrix) and non-osteogenic (stiff matrix) branches, and non-adipogenic (soft matrix) and osteogenic (stiff matrix) branches with support nodes defined (Fig. S4a). Pseudotime trajectories of ECM remodeling and cell adhesion genes that were highly expressed in naïve state remained upregulated during matrix conditioning on stiff matrices (Fig. 3c-i,ii). Soft-matrix adipogenic and non-differentiated branches were characterized by upregulation of adipogenic gene markers and endochondral ossification gene markers, respectively (Fig. 3c-i, top, red and green). The expression trajectory of the master regulator of adipogenesis *CEBPA* monotonically increased along the SoftAD branch and was retarded along the StiffND branch (Fig. S4b-i). Suppression of *CEBPA* expression by stiffness cues further attenuated expression of downstream adipogenic markers including *G0S2*, *LPL*, and *ADIPOQ*. We detected upregulation of endochondral ossification gene markers during matrix conditioning of non-adipogenic cells on soft but not on stiff matrix (Fig. S4b-ii). The stiff-matrix osteogenic branch was characterized by upregulation of osteogenic gene markers (Fig. 3c-ii, blue), which paralleled activation of SRF target genes that mediate mechanical cues and direct cell differentiation (Fig. S4c).^53,54^

### Identification of matrix-responsive genes

Bifurcation of the diffusion pseudotime maps highlights matrix-directed adipogenic and osteogenic differentiation by fat- and osteoid-like elasticities. Matrix-responsive genes are thus characterized by the divergence of their pseudotime expression trajectories between SoftAD and StiffOS branches (Fig. 4a). To identify genes that were statistically significantly responsive to matrix, we first discarded genes with permutation test *p*-values greater than 0.01. The remaining genes were scored for matrix-responsiveness (*MR*) by the area enclosed between SoftAD and StiffOS intensity profiles normalized by mean intensities:

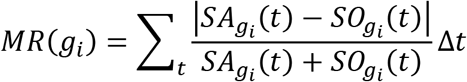

**Figure 4.**
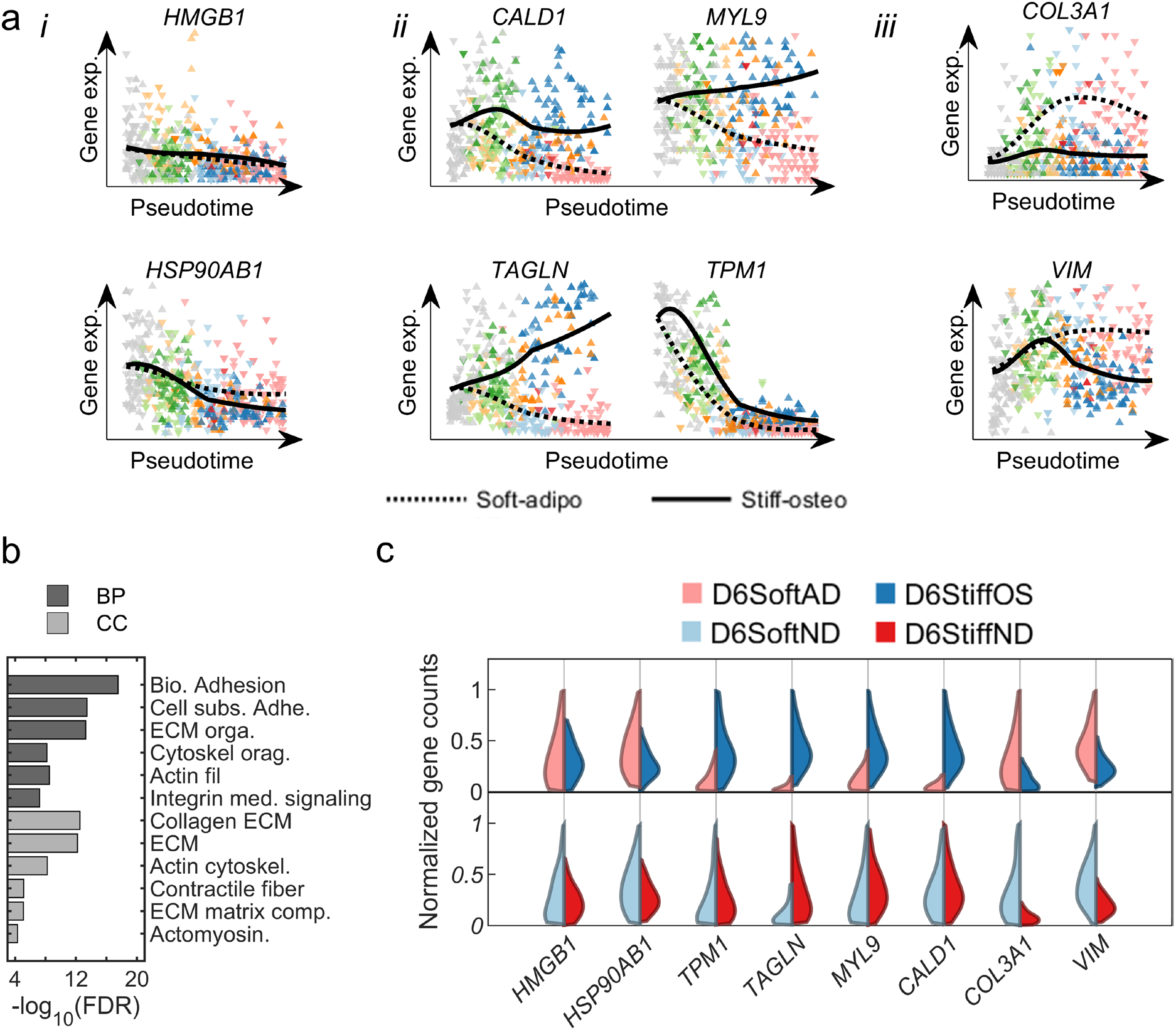
Matrix-responsive genes identified using pseudotime trajectories: **a,** Pseudotime expression trajectories are shown for (i) representative housekeeping genes and (ii) highly ranked matrix-responsive genes that are upregulated by matrix stiffness and (iii) highly ranked matrix-responsive genes that are upregulated by matrix softness. **b,** The GO terms enriched in the top 100 matrix-responsive genes. **c,** Single-cell distributions of upregulation of cytoskeletal genes by matrix stiffness in matrix-directed differentiating subpopulations (top) and non-differentiating subpopulations (bottom). BP: Biological process. CC: Cellular component.

Here, *SA*_*gi*_(*t*) and *SO*_*gi*_(*t*) are the intensities of gene *g*_*i*_ along the SoftAD and StiffOS projections, respectively. The top-ranked genes were enriched for those involved in ECM remodeling, matrix adhesion, and actomyosin cytoskeletal organization (Fig. 4b).

Unlike housekeeping genes *HSP90AB1* and *HMGB1*, which showed no matrix dependence (Fig. 4a-i), genes that were upregulated on stiff matrices both during matrix conditioning and during early differentiation are enriched for actin-binding cytoskeletal components that belong to the so-called cellular contractome^55^ and that are regulated by the SRF mechanotransduction signaling pathway.^56^ The identified genes include *CALD1* and *MYL9*, which encode proteins that regulate myosin head ATPase activity, tropomyosin-1 (*TPM1*, ranked 19), which regulate actin-myosin interactions, and transgelin, which is an actin cross-linker (Fig. 4a-ii). *THY1* is also upregulated on stiff matrices; the protein it encodes directs osteogenesis.^57,58^ *COL3A1*, which is a marker of endochondral ossification, and *VIM*, which encodes a type-III intermediate filament that is expressed in mesenchymal cells and contributes to adipogenesis,^59^ are both upregulated on soft matrices, (Fig. 4a-iii).

### Tropomyosin-1 mediates matrix-directed cell-fate decisions

The expression of the top-scored matrix-responsive genes that encode proteins involved in actin binding is more sensitive to matrix stiffness across progenitor cells that have the capacity to differentiate toward fat and bone (D6SoftAD and D6StiffOS, Fig. 4c, top) than in cells that exhibit impaired matrix-directed differentiation (D6SoftND and D6StiffND, Fig. 4c, bottom). *TPM1*, which is ranked 19, is of particular interest, as it encodes a protein that regulates myosin contractility on soft matrices.^60^ To study the regulation of *TPM1* by matrix elasticity, we performed quantitative immunofluorescence using an antibody that recognizes TPM1 and TPM2 isoforms. *TPM1* was upregulated both at the RNA level (Fig. 4c) and at the protein level on stiff matrices compared to soft matrices concomitantly with increased F-actin polymerization (Figs. 5a-i and 5a-ii). However, TPM1 expression was more sensitive to matrix elasticity than F-actin polymerization (Fig. 5a-iii).

**Figure 5.**
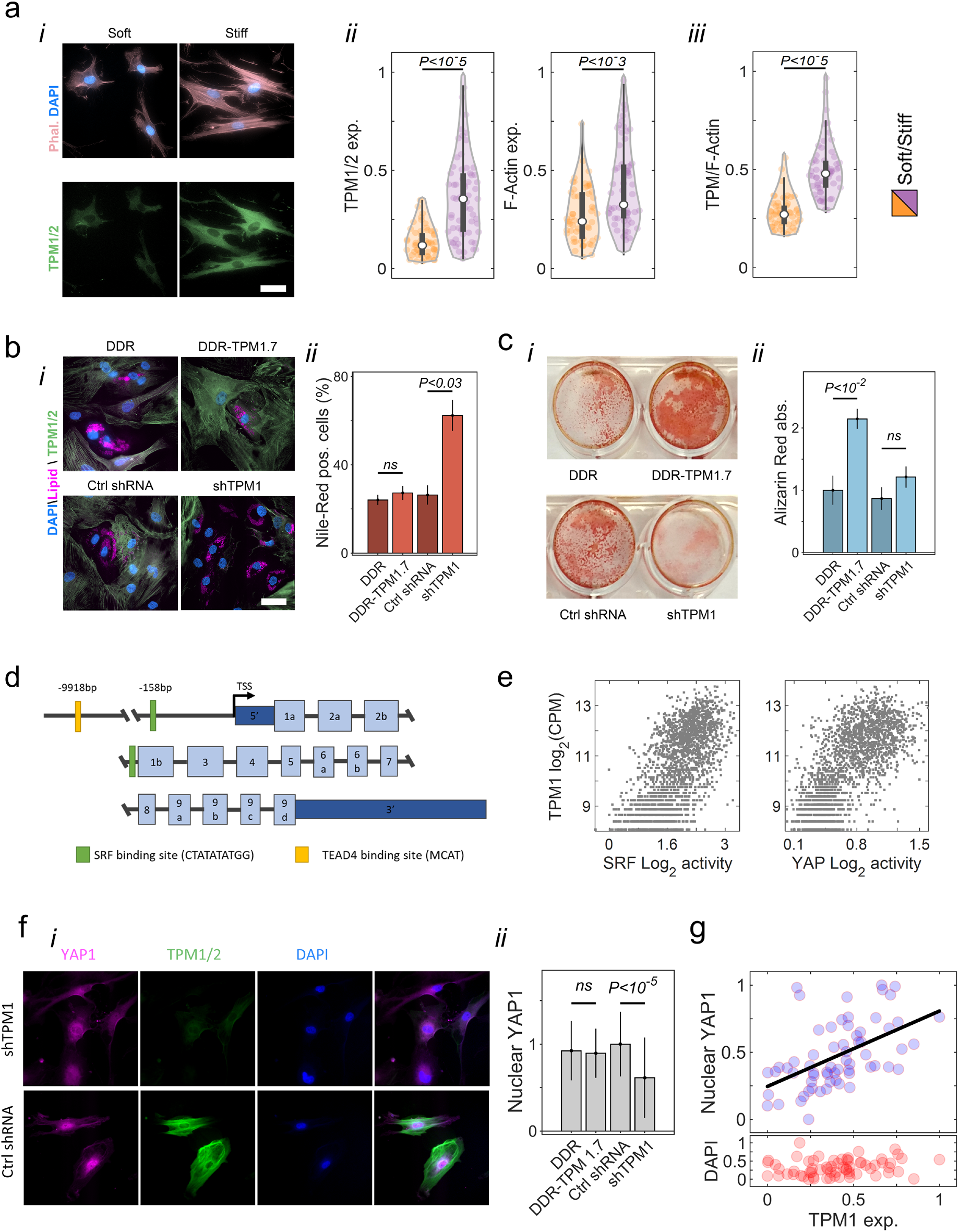
TPM1 is mediates matrix-directed cell differentiation. **a,** (i) Immunofluorescence staining of matrix-conditioned MSCs (Day-3) shows upregulation of tropomyosin expression on stiff matrices concomitantly with F-actin polymerization. Scale bar: 50 μm. (ii) Quantitative analysis of immunofluorescence of tropomyosin (left) and F-actin polymerization (right) on stiff matrices. (iii) The distribution of single-cell ratios between tropomyosin expression and F-actin polymerization on soft and stiff matrix. **b**, (i) Images of cells that express DDR, DDR-TPM1.7, a control shRNA, or shTPM1 stained with Nile red staining for lipids. Scale bar: 50 μm. (ii) Quantification of the percentage of Nile red-stained cells demonstrates enhanced adipogenic differentiation when *TPM1* expression is knocked down. **c**, (i) Images of cells that express DDR, DDR-TPM1.7, a control shRNA, or shTPM1 stained with Alizarin red. (ii) Quantification of the percentage of Alizarin red-stained cells demonstrates enhanced osteogenic differentiation in cells overexpressing TPM1.7. **d,** Schematic representation of *TPM1* genomic sequence. **e,** Correlations of *TPM1* RNA levels with SRF (left, Day 6 Spearman coefficient of correlation 0.62) and YAP1 (right, Day 6 Spearman coefficient of correlation 0.31) signaling activities. **f,** (i) Immunofluorescence staining shows decreased nuclear localization of YAP1 in cells that express shTPM1 compared with cells that express control shRNA. Scale bar: 50 μm. (ii) Quantitative immunofluorescence analysis of nuclear YAP1 in response to overexpression of TPM1.7 isoform and knockdown of TPM1 compared with controls (n=39-96 cells). **g,** Nuclear localization of YAP1 increases with tropomyosin levels. (n=70 wildtype cells). Error bars indicate STD.

To study how TPM1 is involved in regulation of cell-fate decisions, we designed lentiviral constructs encoding Dendra2 (DDR) conjugated to tropomyosin-1.7 isoform cDNA sequence (DDR-TPM1.7) and an shRNA complementary to *TPM1* (shTPM1) under control of a puromycin resistance selection sequence (Fig. S5a). We also generated the respective DDR and non-hairpin shRNA control constructs. Expression of the DDR-TPM1.7 construct resulted in an average twofold overexpression, whereas the expression of shTPM1 decreased Tropomyosin-1 levels twofold compared with controls (Fig. S5b). Transduced cells were cultured in adipogenic and in osteogenic induction media, and cell differentiation toward fat and bone lineages was assessed using Nile red and Alizarin red staining. Strikingly, we found that TPM1 knockdown by shTPM1 increased adipogenesis 2.5 fold while TPM1.7 overexpression had no effect (Fig. 5b). Similarly, DDR-TPM1.7 overexpression increased osteogenesis twofold as assessed based on calcium deposits, whereas *TPM1* knockdown had no effect (Fig. 5c).

The promoter-enhancer region of *TPM1* contains an MCAT binding element of TEAD1-to-4 (transcription enhancer factors for YAP/TAZ)^61^ and a CArG box that is recognized by SRF (Fig. 5d).^62,63^ Single-cell analysis revealed a significant positive correlation between *TPM1* expression and the expression of YAP1 and the SRF target genes (Fig. 5e-i,ii).^26,64^ These data implicate TPM1 regulation by SRF and YAP1 downstream of matrix elasticity. To test whether tropomyosin-1 also activates YAP1 transcriptional regulator, we stained YAP1 in TPM1-kncoked down cells compared with cells expressing a non-hairpin shRNA control (Fig. 5f-i). Indeed, a twofold knockdown of TPM1 led to a 30% delocalization of YAP1 from the nucleus (Fig. 5f-ii).^26^ This conclusion is also supported by non-perturbed primary cells, in which nuclear YAP1 protein levels correlated with tropomyosin-1 protein levels (Fig. 5g). We conclude that tropomyosin-1 is a potent cytoskeletal mechanostat that responds to an increase in contractile forces on stiff matrices by strengthening actin cytoskeletal filaments and acting downstream of SRF to mediate matrix signals that direct MSC differentiation toward soft and stiff tissue lineages.

## Discussion

Single-cell transcriptional profiling provides a means to interrogate lineage specification of different cell types during development,^65,66^ tissue regeneration,^67^ tumorigenesis,^68^ immune response,^69,70^ and cell differentiation.^41^ Here, we employed single-cell RNA sequencing for studying the heterogeneity of cell states within a population of bone-marrow derived MSCs. The heterogeneity of primary MSCs, which has a profound impact on their clinical utility,^7^ integrates multiscale contributions of variation between donors, tissues of origin, clones, and single cells (as reviewed previously^4^). From single-cell analysis of cells cultured using the well-established matrix-directed conditioning and differentiation methodology, we identified nine cell subpopulations using unsupervised clustering. MSC subpopulations are characterized by distinctive properties that are related to cell mechanosensitivity and differentiation capacity. As expected, matrix softness supports adipocyte differentiation, and matrix stiffness promotes osteoblast differentiation. However, we also identified and characterized subpopulations of cells that failed to undergo osteogenesis or to upregulate adipogenic markers on stiff matrix and subpopulations of cells that failed to undergo adipogenesis or to upregulate osteogenic markers on soft matrix. This heterogeneity reflects a continuous phenotypic spectrum of bone marrow-derived stromal cells that lie between multipotent mechanosensitive progenitors that differentiate in tune with matrix cues, osteoblastic-committed cells but no adipocytic-committed cells that differentiate independent of matrix cues and cells that lack the differentiation capacity to either lineage (Fig. S1). In addition, we identified a subpopulation of MSCs that were cultured on soft matrices and expressed endochondral ossification programs (Fig. 2c and 3c).

Dynamic mapping of the cell-fate decision processes provides means for screening matrix-responsive genes based on pseudotime trajectories. Of the average 3,800 expressed genes per cell, one of the most strongly matrix dependent genes was *TPM1*, which encodes twelve alternatively spliced isoforms that form coiled-coil parallel dimers and co-polymerize head-to-tail along actin filaments thus regulating the interactions with myosin motors and actin-binding proteins.^71^ Via these interactions, tropomyosin decrease the mechanical flexibility of actin filaments^72,73^ and stiffens the cell cortex,^74^ thus rendering mechanical strength under increasing load. We show here that TPM1 is a cytoskeletal mechanostat with RNA and protein levels highly sensitive to matrix elasticity. Also, we demonstrated that tropomyosin-1 is a remarkably potent regulator of cell differentiation toward fat and bone on soft and stiff matrices, respectively.

Mechanistically, our results indicate that *TPM1* is transcriptionally regulated by the SRF and by other pathways that mediate mechanical cues via YAP1 as facilitated by the CArG box and MCAT binding motifs, which is consistent with previous reports.^54,75–77^ Tropomyosin-actin co-polymerization protects stress fiber integrity and permits high-frequency myosin power-stroke cycles on stiff matrices.^60^ Myosin contractility and the assembly of stress fibers in turn drive nuclear localization of YAP1 and the transcriptional activation of TEAD-target genes,^26,28^ including TPM1.^77^ Tropomyosin-1 was also shown to antagonize the inhibitory effects of the actin-severing proteins Cofilin and Gelsolin that negatively regulate YAP1 in mechanically-relaxed cells^78^ by inducing a non-favorable conformational changes to filamentous actin (F-actin).^79,80^ By suppressing F-actin disassembly and stabilizing a contractile state, tropomyosin mediates stiff matrix signals and directs osteoblast differentiation downstream of YAP1.

## Methods

### Cell harvesting, culture, and differentiation

Bone marrow aspirates were collected from the iliac crest of healthy human donors for allogeneic transplantation under written consent and the approval of the local institutional Helsinki Committee (0626-15-HMO). Aspirates were passed through a nylon cell strainer, separated by a Ficoll-Hypaque density gradient, and resuspended in low-glucose Dulbecco’s Modified Eagle Medium (DMEM; Biological Industries) supplemented with 1% L-glutamine (Biological Industries), 1% penicillin-streptomycin (Biological Industries), and 10% fetal bovine serum (Biological Industries). Cells were seeded into 75-cm^2^ culture flasks (30 × 10^6^ cells per flask) and cultured at 37 °C in a humidified atmosphere with 5% CO_2_. Medium was replaced twice weekly, and cell density was maintained < 80-85% confluence. The positive (CD73 and CD90) and negative (HLA-DR, CD56, CD3, and CD45) surface-marker repertoire was validated by FACS using targeting antibodies (eBioscience).

Adipogenic induction was performed in low-glucose DMEM medium supplemented with 10 μg mL^−1^ insulin, 500 mM 3-isobutyl-1-methylxanthine (Sigma-Aldrich), 1 μM dextran (Sigma-Aldrich), and 100 μM indomethacin (Sigma-Aldrich). Osteogenic induction was performed in low-glucose DMEM medium supplemented with 50 μg mL^−1^ L-ascorbic acid 2-phosphate (Sigma-Aldrich), 10 mM glycerol 2-phosphate (Sigma-Aldrich), and 10 nM dexamethasone (Sigma-Aldrich). Bi-potential induction medium was prepared by mixing adipogenic and osteogenic induction media at equal volumes as reported.^35,36^

### Hydrogels with controlled elasticity

Cells were seeded at 5,000 cells/cm^2^ density onto soft substrates with controlled elasticities (2 kPa and 25 kPa) composed of thin polyacrylamide hydrogel films and uniformly coated with rat-tail type-I collagen (Petrisoft; Matrigen). The hydrogels were coated with a 0.02 mg mL^−1^ rat tail type I collagen solution with a constant surface density.

### Droplet-based single-cell RNA barcoding and library preparation

inDrop high-throughput single-cell labeling was performed as reported with minor adaptations.^39,81^ Briefly, cells were trypsin-detached, washed in phosphate-buffered saline (PBS, Sigma-Aldrich) and maintained dissociated in suspension supplemented with 15% OptiPrep (Sigma-Aldrich). Cells (~1000 cells/μl) were loaded into a chilled syringe pump and injected into a custom-designed microfluidics device. Flow settings were adjusted to allow individual cells to become encapsulated with single hydrogel beads presenting cell-specific barcoding oligonucleotide primers inside 3-5 nL encapsulation mixture droplets that contain 20 U μL^−1^ reverse transcriptase (SuperScript III, Thermo Fisher Scientific), First-Strand Buffer (Thermo Fisher Scientific), 0.6% (v/v) IGEPAL CA-630 (Sigma-Aldrich), 1 mM dNTPs (New England Biolabs), 6.67 mM DTT (Thermo Fisher Scientific), 2700 units mL^−1^ murine RNase inhibitor (New England Biolabs), and 0.1 M Tris-HCl (pH 8.0)).^81^ Primers consisted of a poly-T sequence to trap mRNA molecules, two 8-base cell-specific barcodes, 5-base unique molecular identifier (UMI) sequence, and a T7 promoter sequence. Droplets were collected, the primers were photo-cleaved by exposing the emulsion to UV (365 nm 10 mW cm^−2^, BlackRay Xenon lamp) for 10 minutes at room temperature, and reverse transcription of trapped mRNA was performed at 50 °C for 2 hours. Droplets were then demulsified by adding 1H,1H,2H,2H-perfluoro-1-octanol (Sigma Aldrich) dissolved in HFE-7500 (Novec) at 20% (v/v). Single-stranded DNA primers and primer dimers were digested using 1 unit μL^−1^ ExoI (New England Biolabs) and 1 U μL^−1^ HinFI (Thermo Fisher Scientific). mRNA:cDNA hybrids were purified using Agencout AMPure XP beads (Beckman Coulter Genomics) according to the manufacturer’s instructions. Second-strand cDNA synthesis was performed using NEBNext Ultra II Non-Directional RNA Second Strand Synthesis Module (New England Biolabs) and amplified by *in vitro* transcription (HiScribe, New England Biolabs). RNA products were fragmented using Ambion RNA Fragmentation Reagents (AM8740, Thermo Fischer Scientific), purified using Agencout AMPure XP beads, and reverse-transcribed using random hexamer primers (SuperScript III, Thermo Fisher Scientific). Single-stranded DNA fragments were linearly amplified via 14 PCR cycles (HiFi HotStart ReadyMix, Kapa Biosystems).

### Sequencing and processing of single-cell RNA data

Libraries were paired-end sequenced (PE5: 45 bp and PE7: 35 bp) on a NextSeq 500 platform using a High Output kit (75 cycles, Illumina). Sequences were demultiplexed using indices and barcodes, permitting ≤ 1 nucleotide alignment substitution. After removing low-complexity poly-adenylated sequences, transcripts were aligned against an indexed reference genome (GRCh38.p12) using Bowtie 0.12.8, and gene intensities were evaluated based on UMI frequency. Cells with high representation of mitochondrial RNA (>25%) or low gene number (≤1,900 per 15,000 UMIs) were discarded. Mitochondrial and ribosomal genes were discarded as well as low-frequency genes that were detected in ≤ 5 cells. Finally, UMI count was down-sampled to 15,000 per cell, thus normalizing gene expression and minimizing batch-specific confounders.

### Clustering MSC subpopulations

Computational and statistical analyses were performed using established protocols in R (R Foundation for Statistical Computing) and Matlab (Matlab 2019 Mathworks Ltd). We distinguished highly variable genes from technical noise as described previously by Brennecke and colleagues.^82^ Outlier expression values were truncated by winsorizing, the squared coefficient of variation (CV^2^) versus the mean, *μ*, was fitted by *α/μ* + *β* (**Error! Reference source not found.**), and highly variable genes were identified using the chi-squared test (FDR < 10^−3^). UMI counts (CPM) of highly variable genes were log_2_(*x* + 1) transformed and single-cell transcriptomes were dimension-reduced via PCA. The number of significant PCs was determined based on a screen test. Unsupervised cell clustering was performed using *k*-means clustering in the PCA space. The number of clusters was optimized based on within-cluster sum of squares. Genes that were differentially expressed across clusters were assigned based on the corresponding log-fold ratios of expression levels and the adjusted two-tailed Wilcoxon rank-sum test *p*-values (Benjamini–Hochberg method). Functional enrichment analysis was performed using Gene Set Enrichment Analysis database.^83^

### Scoring adipogenic-osteogenic differentiation

Adipogenic and osteogenic differentiation was scored based on a subset of established markers (adipocytes: *PPARG*, *CEBPA*, *PLIN1*, *G0S2*; osteocytes: *COMP*, *TMEM119*, *DCN*, *BCAP31*, *ITGA5*, *COL1A1*, *ATF4*, *CREB3L1*, *FSTL3*) using:

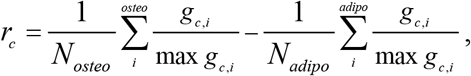

where *g*_*c,i*_ is the transcript level of gene *i* in cell *c*.

### Scoring oxidative phosphorylation state

Oxidative phosphorylation was scoredusing a set of 23 genes that encode proteins of the electron transport chain (complexes I-V) and the ATP synapse complex. The single-cell expression intensities of these genes were z-score normalized and averaged. D3soft subpopulation was divided into low oxidative (< −0.23), mid (> −0.23 and > 0.13) and high (> 0.13) oxidative phosphorylation cells.

### Diffusion pseudotime analysis

Diffusion mapping was performed using the Destiny package in R.^84^ Only genes that varied significantly across soft and stiff matrix-conditioned cells, which we identified using the CV^2^ plot as described above (FDR < 10^−3^), were retained. Diffusion mapping was calculated separately for cells that were cultured on soft matrices (NV, D3Soft and D6Soft; Fig. 3a) and cells that were cultured on stiff matrices (NV, D3Stiff and D6Stiff; Fig. 3b). For each matrix, we performed PCA dimensionality reduction, and the diffusion components were calculated based on the top twenty principle components that maximized variance between transcriptomes with the local sigma calculated by nearest-neighbour approximation (k = 50).

Matrix-directed cell differentiation dynamics was analysed using the MeRLOT package in R.^85^ Soft and stiff matrix scaffold trees were reconstructed using the first two diffusion components (CalculateScaffoldTree function in R). Principle elastic trees were calculated, and cells were assigned to 50 discrete support nodes that were aligned along the bifurcating branches (CalculateElasticTree function in R). Setting the first support nodes that are associated with naïve MSC state as the initial pseudotime, pseudotime values were further propagated along the bifurcating branches of the support trees in proportion with the distance from *t*_0_ and cells were assigned accordingly (CalculatePseudotimes function in R). Next, cells were binned into pseudotime sets (25 bins for the trunk and for each of the branches), and dynamic gene expression profiles were averaged within each bin along the bifurcating trajectories of the soft and stiff matrix trees.^86^ Gene trajectories were further smoothed by fitting a natural cubic spline with three degrees of freedom (Figs. 4a and Fig. S4b).

Gene trajectories along the Soft-Adipo and Stiff-Osteo were clustered using *k*-means, and the clusters were chronologically ordered according to the average pseudotime point of maximal gene expression. Genes that were differentially expressed between cell subpopulations (lfc > 1, adjusted FDR < 10^−5^) were ranked by similarity using a cosine-distance metric within each cluster, and trajectories of most similar genes within each cluster were plotted with respect to SoftAD (Fig. 3b-i) and StiffOS (Fig. 3b-ii) branches.

### Permutation test of mechanically diverging genes

The statistical significance of gene divergence along the SoftAD versus StiffOS trajectories was evaluated using a permutation test. Specifically, cells were randomly reassigned a branch label while maintaining the correct overall distribution of cells between branches. New branch curves were then calculated, and the normalized divergence area between the curves was calculated as described above. A distribution of the divergence area was obtained based on 1,000 random permutations, and *p*-value of the non-permutated divergence was calculated.

### Pathway analysis

The activation of SRF and YAP1 signaling was evaluated by measuring the expression levels of respective target genes.^87,88^ Single-cell expression levels of target genes showing a minimum average expression (UMI > 0.5) were linearly rescaled to [0,1], and signaling pathway activation was scored in each cell by the average normalized intensity.

### Sample preparation and optical imaging

Following culture, cell samples were washed with PBS, fixed in 4% paraformaldehyde, blocked in 2% BSA with 0.5% (v/v) Triton X-100 in PBS for 20 minutes, immersed in 2% BSA for 1 hour, and rinsed in PBS. Staining using the primary anti-TPM1/2 antibody (TM311, Sigma-Aldrich) and anti-YAP1 antibody (Proteintech), and the secondary donkey anti-mouse IgG antibody (Alexa Fluor 647; 1:100; Abcam), donkey anti-rabbit IgG antibody (Alexa Fluor 647; 1:100; Abcam) and donkey anti-mouse IgG antibody (Alexa Fluor 488; 1:100; Abcam) was performed according to manufacturer’s protocols. DAPI (Sigma-Aldrich, 20 min) and Phalloidin-iFluor 555 (Abcam) staining (165 nM, 30 minute immersion) was performed in PBS. Lipid droplets staining was performed by immersing cells in 0.1 μg mL^−1^ Nile red (Sigma) for 5 minutes. Immunofluorescence imaging was performed using a NIKON Ti-E inverted microscope equipped with an sCMOS iXon3 camera (Anodr) and a Spectra X light engine light source (Lumencor). A CFI Apo TIRF 60X Oil (Nikon) and a CFI Plan Apo VC 20X (Nikon) objectives were used. Cell and nucleus projected areas were segmented and quantified using custom-built MATLAB code.

### Calcium deposit quantification

MSC calcium deposits were stained using Alizarin red according to established protocols. Alizarin red was dissolved in doubly distilled water (2% w/v), and the pH was adjusted to 4.2 using HCl. Fixed samples were rinsed in PBS, immersed in Alizarin red solution for 15 minutes at room temperature and rinsed twice with doubly distilled water. Alizarin red S-calcium complexes were extracted by immersing in 0.5 N HCL/5% SDS (w/v) extraction solution. The concentration of the extracted stain was quantified by measuring the absorbance at 410 nm (SmartSpec 3000 spectrophotometer, Bio-Rad) and normalizing by the number of cells in each sample as measured via Hoechst 33342 (Thermo Fisher Scientific) staining.

### TPM1 knockdown and overexpression

Knockdown of *TPM1* was performed using the pLKO.1 plasmid lentiviral backbone (a kind gift from Bob Weinberg, Addgene plasmid #8453) either encoding an shRNA with sequence complementary to *TPM1* (shTPM1: 5’-CGGAGAGGTCAGTAACTAAAT-3’) or a control non-hairpin insert (5’-CCGCAGGTATGCACGCGT-3’). Selection of expressing cells was performed in the presence of 1 mg ml^−1^ puromycin (Sigma-Aldrich) for 2 weeks following viral infection and confirmed using mCherry fluorescence signal. TPM1.7 overexpression was performed by generating a human MSC-derived cDNA library. The *TPM1.7* sequence was amplified using dedicated primers, a sequence encoding a C-terminal DDR sequence was conjugated, and the fragment was cloned into a lenti-EFIα pEIGW expression vector. Control overexpression vector did not contain the *TPM1.7* sequence.

Lentiviral particles were generated using human embryonic kidney (HEK) 293T cells. HEK cells were seeded in 55 cm^2^ plates at 50% confluence. Transfer (10 μg), packaging (10 μg psPAX2, Addgene #12260), and envelope (6 μg pMD2.G, Addgene #12259) viral plasmids were diluted in 500 μL serum-free DMEM. Next, plasmids were mixed with 500 μL polyethylenimine (PEI, Sigma-Aldrich) dissolved in DMEM for a final ratio of 1:2.5 DNA to PEI. HEK cells were co-transfected by incubating with DNA-PEI complexes for 18 hours. Medium was exchanged and supernatant was collected after 24 and 48 hours, filtered (0.45 μm PVDF, Millex), and MSCs (passage 1-2) were infected and transduced with lentiviral particles.

## Supplementary Figures

**Figure S1.**
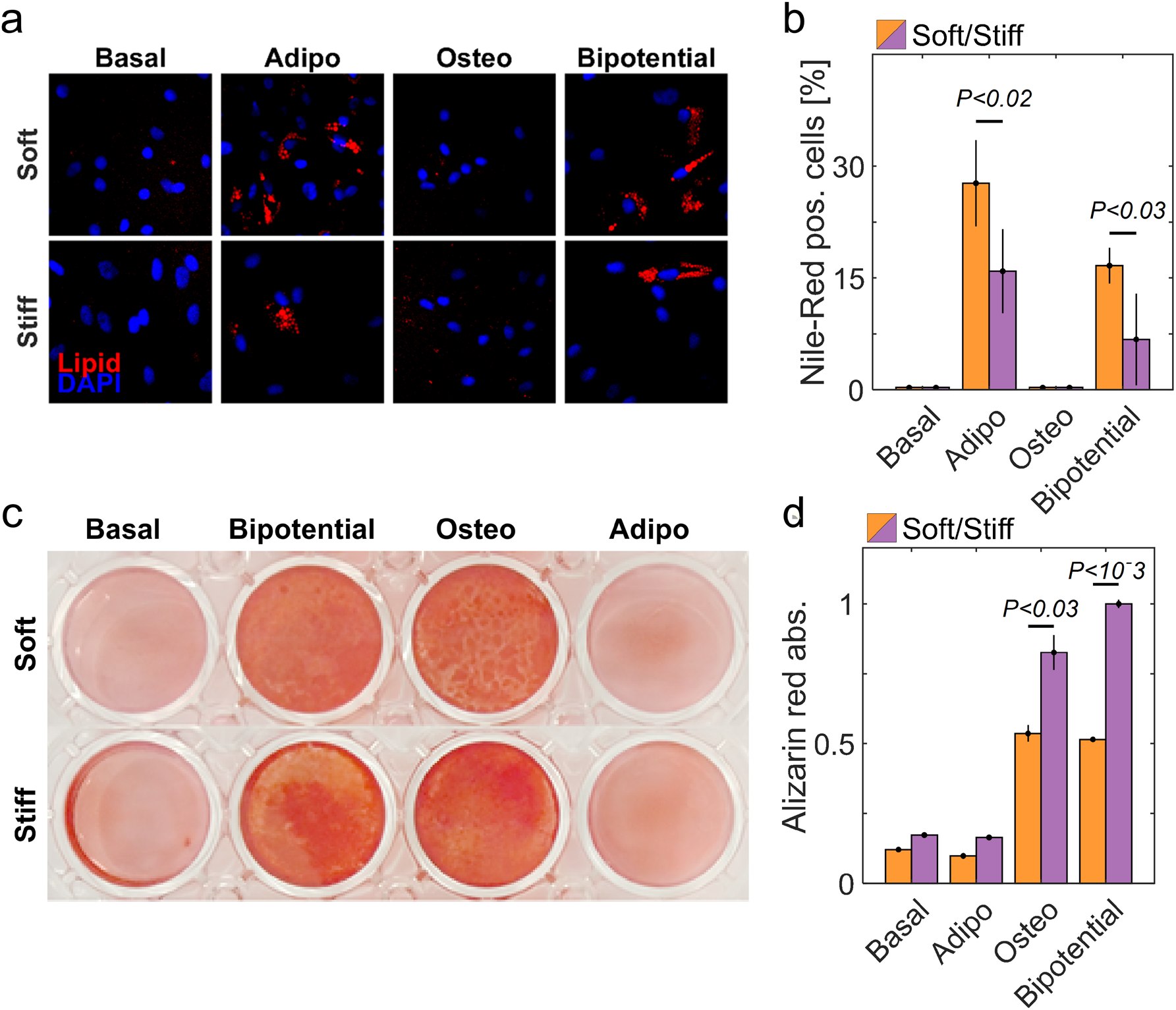
Matrix elasticity directs MSC differentiation toward soft and stiff tissue lineages. **a,** Nile red lipid staining marks adipogenic differentiation of cells cultured on soft and stiff matrices for 10 days in basal medium and cells cultured for 3 days in basal medium followed by 7 days in adipogenic, osteogenic, or bi-potential induction media. **b,** The fractions of positively stained cells show upregulation of adipocyte differentiation by soft matrices only under supporting medium conditions. **c,** Alizarin red staining of calcification marks osteogenic differentiation of MSCs cultured for 17 days in basal medium and cells cultured for 3 days in basal medium followed by 14 days in adipogenic, osteogenic, or bi-potential induction media. **d,** Average Alizarin red intensities show basal osteogenesis under all medium conditions and upregulation by stiff matrices under supporting medium conditions. Error bars indicate STD.

**Figure S2.**
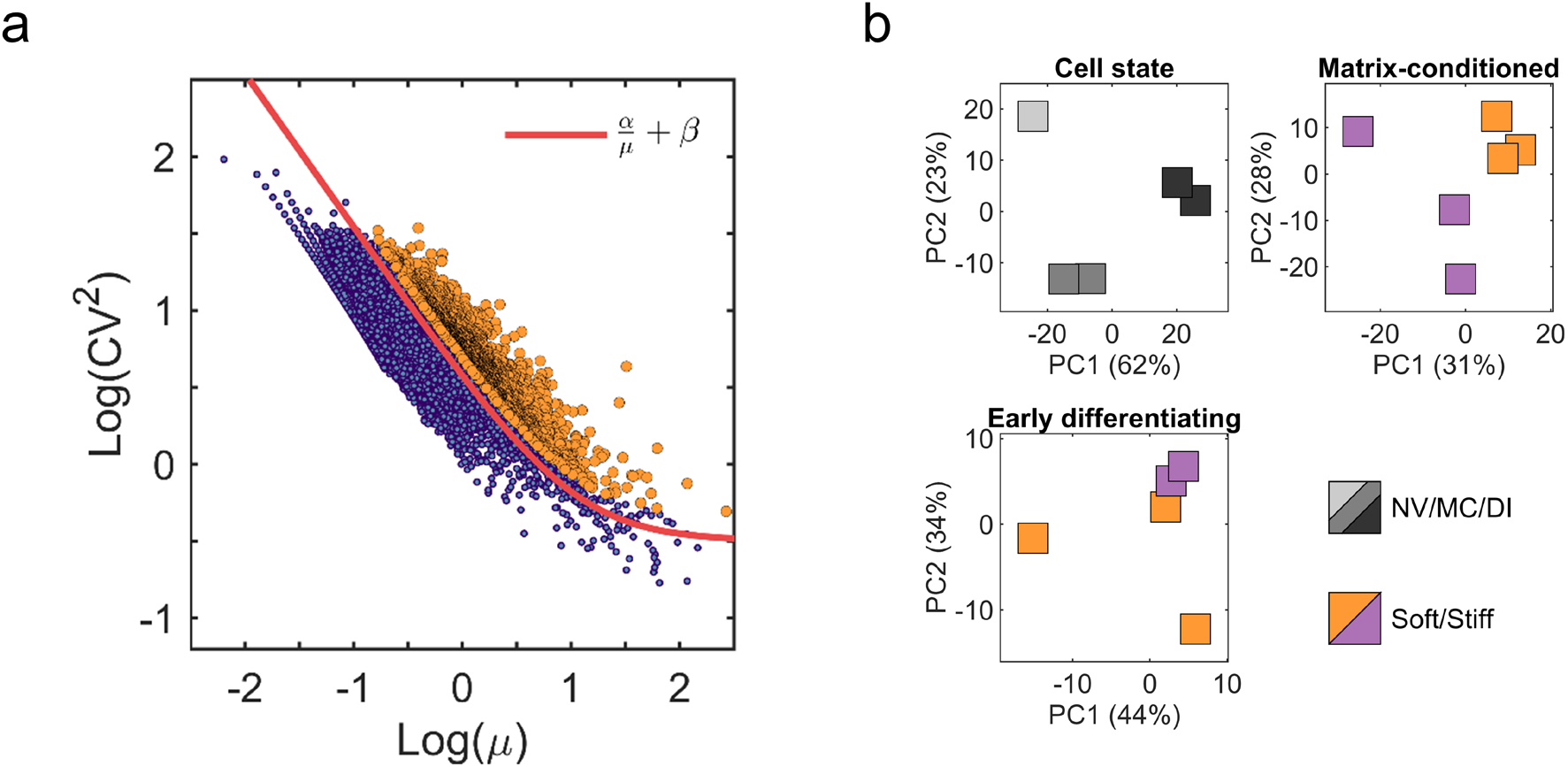
Single-cell transcriptomes are clustered based on culture settings and matrix elasticities. **a,** Standard CV^2^-mean plot identifies highly variable genes across single-cell transcriptomes. **b,** Single-cell transcriptomes cluster according to cell state (top left). Matrix conditioned (top right) and early differentiating (bottom-left) cells cluster according to matrix elasticity. NV: naïve. MC: matrix conditioned. DI: differentiation induction.

**Figure S3.**
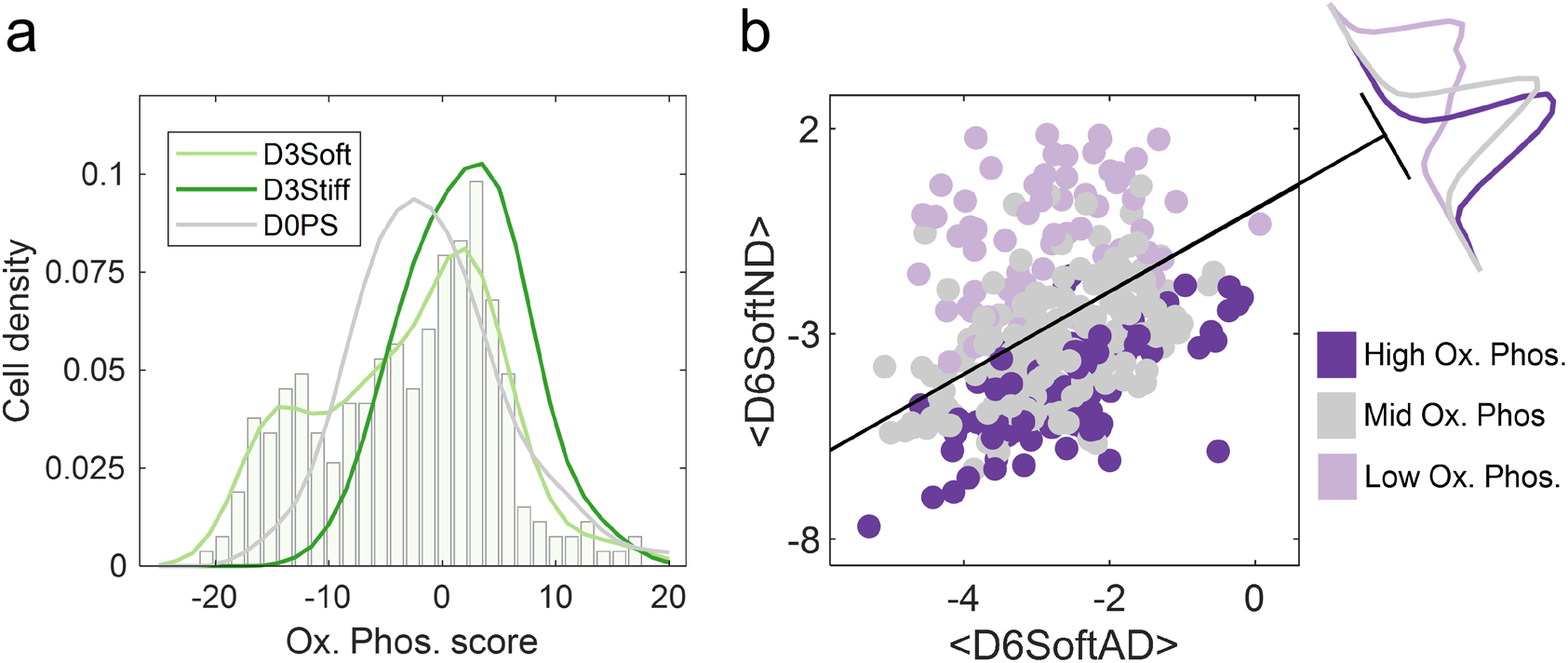
Soft matrix supports low-oxidative phosphorylation cells directed towards endochondral ossification. **a,** Comparison between single cell distributions of naïve D0PS cells and matrix-conditioned D3Soft and D3Stiff subpopulation reveal the emergence of low OxPhos scoring cells only on soft matrix. **b,** Projection analysis onto the transcriptome averages of early differentiating soft-matrix subpopulations calculated relative to differentially expressed genes (flc>1; FDR<10^−5) establish an association of low oxidative-phosphorylation soft matrix conditioned cells with endochondral ossification genes upregulated in D6SoftND cells.

**Figure S4.**
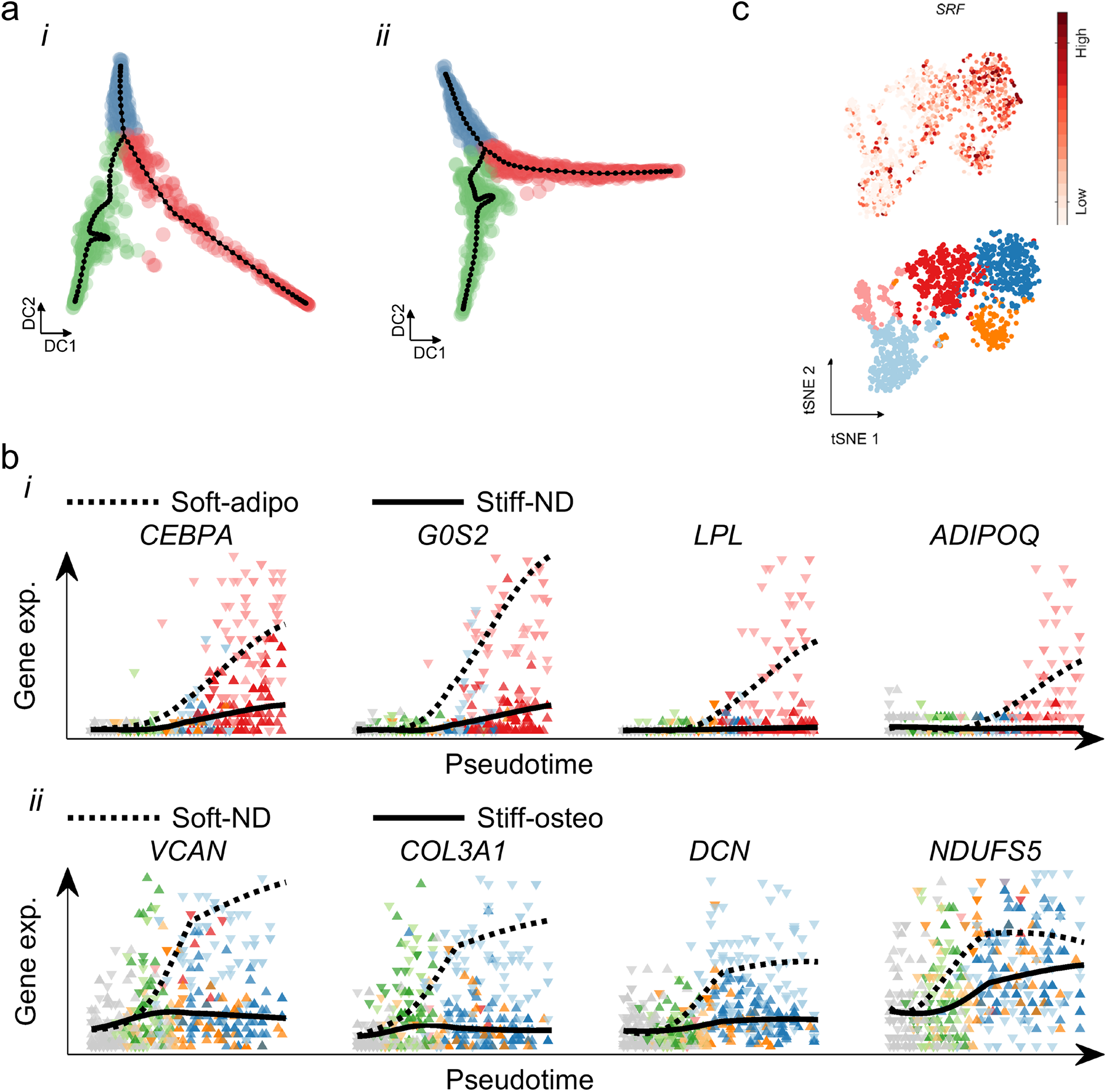
Soft matrices direct adipogenesis of multipotent cells and support endochondral ossification of non-adipogenic cells. **a,** Scaffold tree branch assignments of matrix conditioning (green), adipogenic/non-osteogenic (red), and osteogenic/non-adipogenic (blue) fates on (i) soft and (ii) stiff matrices serve as templates for principle elastic tree support nodes (black). **b,** (i) Pseudotime gene trajectories show upregulation on soft matrix and suppression in non-osteogenic cells cultured on stiff matrix of *CEBPA*, the master adipogenic regulator, and downstream adipogenic markers. (ii) Soft matrix signals support endochondral ossification and suppress electron transport chain gene expression in non-adipogenic cells.

**Figure S5.**
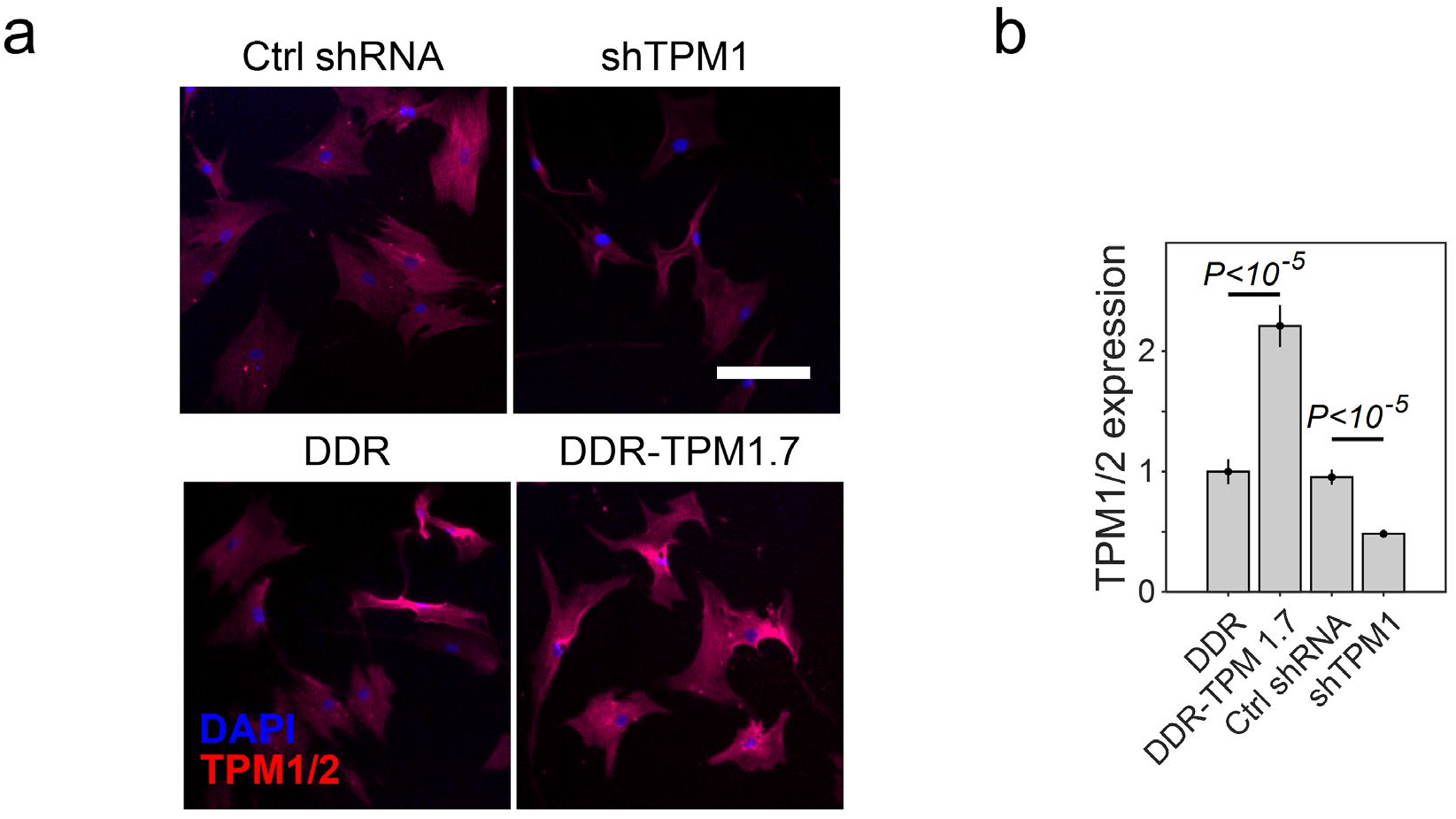
TPM1 knockdown and overexpression. **a,** Images of cells transduced with vectors for expression of shRNA and DDR controls and shTPM1 and DDR-TPM1.7 and stained for tropomyosin isoforms. Scale bar: 100 μm. **b,** Quantitative immunofluorescence analysis of tropomyosin levels shows approximately twofold upregulation in cells overexpressing TPM1.7 and twofold downregulation in cells expressing shTPM1 compared with controls. DDR: Dendra2 control. DDR-TPM1.7: Dendra2-conjugated tropomyosin 1.7. Ctrl shRNA: Non-hairpin insert sequence. shTPM1: shRNA targeting *TPM1*.

## Acknowledgments

AB greatly appreciates support from the U.S.-Israel Binational Science Foundation (BSF 2017357) and Israel Science Foundation (Grant number 1246/14). OR is supported by research grants from the European Research Council (ERC-StG 715260), the Israeli Center of Research Excellence (I-CORE) program, the Israel Science Foundation (Grant number 1618/16), and the Azriely Foundation Scholar Program for Distinguished Junior Faculty. AB and OR thank N. Friedman and A. Ben-Zvi (Hebrew University of Jerusalem), and C. Luxenburg (Tel Aviv University) for fruitful discussions. Author contributions: AB and OR formulated and designed the research program. SB performed all experiments and computational analyses. DB and AM assisted in performing microfluidic-based single-cell RNA sequencing experiments. YKT carried out clustering and classification computations. SB, BA, OR, and AB conducted discussions and brainstorming. BA provided primary mesenchymal stem cells and clinical insight. OR and AB supervised research. SB and AB designed and prepared the graphical material. AB wrote the text. Competing interests: No financial or conflict of interests are declared.

